# Biased belief updating and suboptimal choice in foraging decisions

**DOI:** 10.1101/713941

**Authors:** Neil Garrett, Nathaniel D. Daw

## Abstract

In many choice scenarios, including prey, employment, and mate search, options are not encountered simultaneously and so cannot be directly compared. Deciding which ones optimally to engage, and which to forego, requires developing accurate beliefs about the overall distribution of prospects. However, the role of learning in this process – and how biases due to learning may affect choice – are poorly understood. In three experiments, we adapted a classic prey selection task from foraging theory to examine how individuals kept track of an environment’s reward rate and adjusted their choices in response to its fluctuations. In accord with qualitative predictions from optimal foraging models, participants adjusted their selectivity to the richness of the environment: becoming less selective in poorer environments and increasing acceptance of less profitable options. These preference shifts were observed not just in response to global (between block) manipulations of the offer distributions, but also to local, trial-by-trial offer variation within a block, suggesting an incremental learning rule. Further offering evidence into the learning process, these preference changes were more pronounced when the environment improved compared to when it deteriorated. All these observations were best explained by a trial-by-trial learning model in which participants estimate the overall reward rate, but with upward vs. downward changes controlled by separate learning rates. A failure to adjust expectations sufficiently when an environment becomes worse leads to suboptimal choices: options that are valuable given the environmental conditions are rejected in the false expectation that better options will materialize. These findings offer a previously unappreciated parallel in the serial choice setting of observations of asymmetric updating and resulting biased (often overoptimistic) estimates in other domains.

## Introduction

In contrast to classic economic decisions in which individuals choose from a menu of well-defined options presented simultaneously (De Martino et al., 2013; FitzGerald et al., 2009; Frank et al., 2004; Hunt et al., 2012; Kable and Glimcher, 2007; Rangel and Hare, 2010; Smith et al., 2010; Tom et al., 2007), the options in many real-world decisions are encountered serially and cannot directly be compared to one another. For instance, should I take a currently available option (e.g., hire this candidate for a position or accept this proposition of a romantic date)? Or should I forgo it in the expectation that a better option will come along? Insofar as accepting an option may require passing up unknown opportunities that might arise later, these choices involve opportunity cost and require comparing each prospect to some measure of the overall distribution of alternatives.

Such serial decisions arise canonically in several stylized tasks from animal ethology and optimal foraging theory, such as the prey selection task (Krebs et al., 1977; Stephens and Krebs, 1986). Here, a predator encounters a series of potential prey, which may each be accepted or forgone. If accepted, an option provides gain (e.g., calories) but pursuing and consuming it costs time that could instead be used to search for additional prey. The optimal choice rule, given by the Marginal Value Theorem (MVT, Charnov, 1976), is to accept an option only if its local return (calories divided by time) exceeds the opportunity cost of the time spent. This is just the calories per timestep that otherwise would be expected to be earned on average: the overall, long-run reward rate in the environment. Thus, foragers should be pickier when the environment is richer: a mediocre target (e.g., a skinny, agile animal that takes time to chase down) may be worthwhile in a barren environment, but not in one rich with better alternatives.

In this setting, then, choice turns largely on estimating the environment’s present rate of return, which establishes the threshold against which each prospective prey item can be assessed. Importantly, the MVT specifies the optimal static choice policy in an environment, and thus predicts differences in behavior between environments. But it does not prescribe the dynamics of how an organism might attain the optimum. Accordingly, its predictions have most often been studied in terms of asymptotic behavior in stable environments. But estimating an environment’s richness from experience with offers is in fact a learning problem, especially in dynamic environments in which the availability of resources fluctuates over time, as with the seasons (Constantino and Daw, 2015; Hutchinson et al., 2008; McNamara and Houston, 1985). Here we ask how humans update beliefs about their environment’s rate of return in a prey selection task.

Previously, we have shown (Constantino and Daw, 2015) that when individuals undertake a different type of foraging problem (“patch leaving”), their choices are well explained by an error-driven incremental learning rule for estimating the environment’s reward rate (Schultz et al., 1997; Sutton and Barto, 1998), which then serves as a comparator for determining which prospects are acceptable. However, systematic deviations from optimal thresholds were detected, which are even more pronounced in participants under stress (Lenow et al., 2017) or with depleted levels of dopamine (Constantino et al., 2017). This suggests that there are circumstances under which individuals may misestimate the environment’s rate of return, and in doing so make choices out of step with what the MVT would predict. However, in the line of studies on patch foraging, these biases have simply been treated as fixed choice tendencies, and in particular have not been shown to arise from the underlying learning process.

Compared to patch leaving, prey selection tasks should in principle provide a cleaner window on learning of environmental reward rates. This is particularly because they lack the structured autocorrelation among outcomes that arises due to the depletion of patches over repeated visits and the threshold-based patch-leaving rule, which makes it difficult directly to correlate trial-by-trial outcomes with choices to reveal learning. However, to date prey selection has only been examined in non-human animals, and without an explicit learning component. Here, evidence for performance consistent with the MVT is mixed. For instance, primates (albeit highly trained under relatively stable conditions) closely approximate the MVT (Blanchard and Hayden, 2014). Conversely, in a study involving large reward rate changes between environments, rats strongly deviated from the MVT by failing to reject costly options even in rich environments (Wikenheiser et al., 2013). The authors proposed a model in which this suboptimality arose due to a fixed subjective cost for rejecting options, although again explanations in terms of learning have not so far been examined.

Here we adapt the trial-by-trial learning model previously used for patch foraging (Constantino and Daw, 2015; McNamara and Houston, 1985) to the prey selection task and extend the model to account for specific deviations from the MVT that we observe. To do this, we fit the behavior of participants performing a novel gamified version of the prey selection task. Participants were tasked with choosing whether to accept or reject serially presented options which varied in terms of points earned and time expended (if accepted). By manipulating the rate of reward between blocks, we were able to examine how individuals responded to changes in the overall richness of their environment as well as examine dynamic trial by trial adjustments that occurred within blocks. We fitted participants’ behavioral choices to two separate learning models. The first (Symmetric Model) allowed beliefs about the environment’s reward rate to update using a single learning parameter. The second (Asymmetric Model) learned to a different degree from positive vs. negative prediction errors. This feature, which has been observed in a number of other learning scenarios (Eil and Rao, 2011; Garrett and Sharot, 2014, 2017; Garrett et al., 2014, 2018; Korn et al., 2012; Kuzmanovic and Rigoux, 2017; Kuzmanovic et al., 2015, 2016; Lefebvre et al., 2017; Mobius et al., 2011; Moutsiana et al., 2013, 2015; Sharot and Garrett, 2016; Sharot et al., 2011) but not explored in the context of foraging, gave the model the capacity to express learning asymmetries and to predict and capture specific patterns of experience-dependent biases in choices.

Across three experiments, participants were sensitive to both global and trial-to-trial changes in the environment, in the direction predicted by the MVT (Charnov, 1976; Stephens and Krebs, 1986). But in all experiments, the Asymmetric Model provided a superior fit to participants’ behavior relative to the Symmetric Model, reflecting information integration being greater for positive compared to negative prediction errors. This asymmetry accounted for a key deviation from the MVT-predicted policy: a reluctance to revise beliefs and change choices when environments deteriorated, leading in that circumstance to an overoptimistic bias and a pattern of overselective choices.

## Results

We first conducted two online experiments (see **Fig. 1** and **Methods** for further details). On each trial, participants were offered an option (styled as an alien) and chose whether to accept or reject it. Accepting it provided a reward (in the form of points, later converted to a bonus payment) but also incurred an opportunity cost in the form of a time delay. There were four possible options, comprising of all combinations of low or high reward after low or high delay (low delay, high reward: LDHR, low delay, low reward: LDLR; high delay, high reward: HDHR; high delay, low reward: HDLR; **Fig. 1b**). Following an accept decision, participants had to wait for the delay to elapse before the points were accrued and the next trial began. Following a reject decision, the experiment progressed to the next trial.

**Figure 1.**
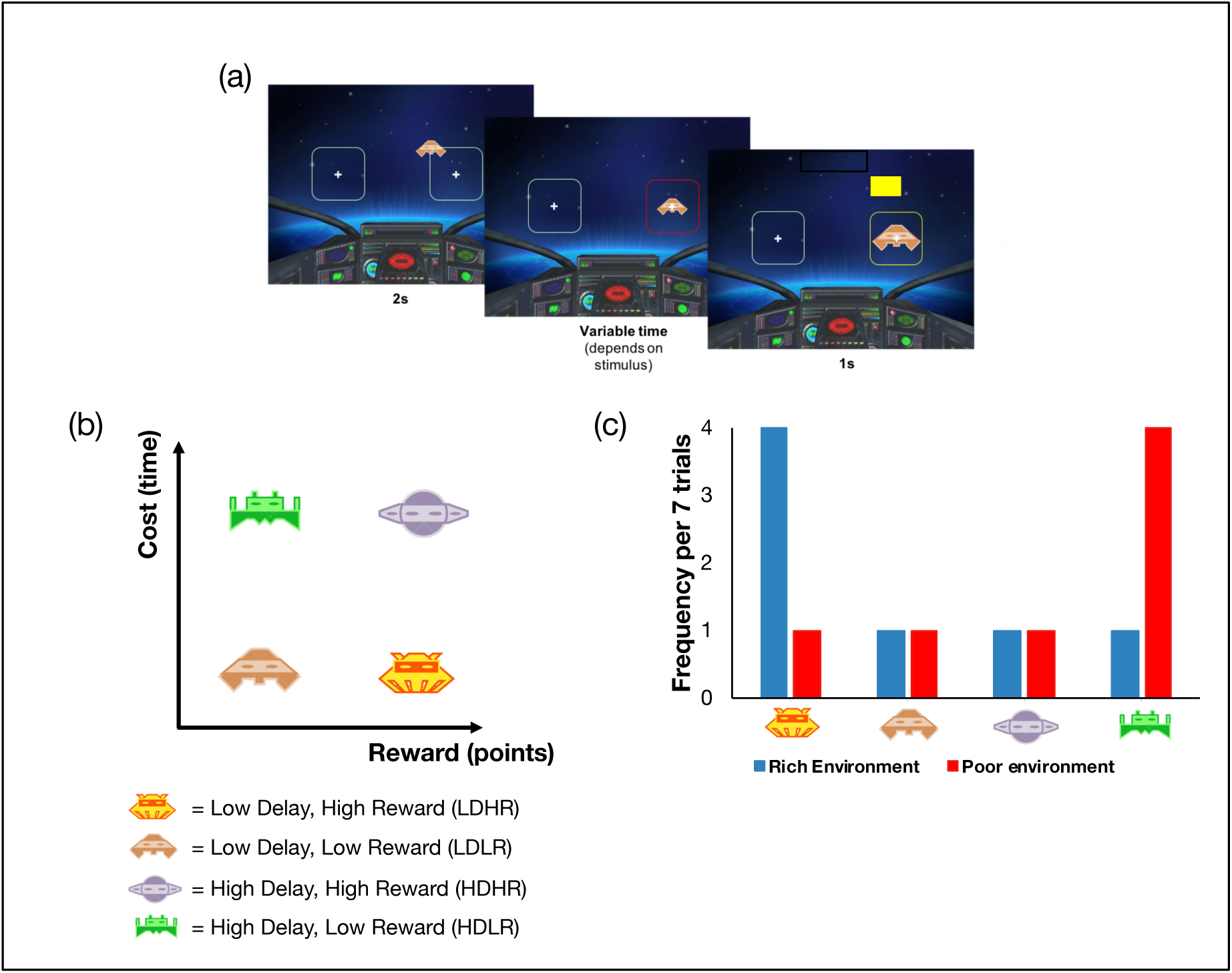
Behavioral task and variables. (a) Timeline of the task. On each trial, an option (presented as one of four different aliens) approached one of two targets. Participants decided to accept the option by selecting the target the option approached (right target in this example) or reject the option by selecting the alternative target (i.e. the target the option did not approach, left target in this example). When options were accepted, the selected target changed color to red and participants were required to maintain a key press during which the option expanded within the target. The target then changed color to yellow indicating that participants could release the key press and the number of points obtained was displayed represented as a partially filled vertical bar. When options were rejected, the experiment immediately progressed to the next trial. (b) On each trial, participants encountered 1 of 4 possible options each of which provided either a low/high reward (points, later converted to a bonus payment) but also incurred either a short/long opportunity cost in the form of a time delay. (c) The experiment was divided into 2 blocks (environments) and the frequency of the 4 different options varied in each of these. There was a rich environment in which the best option (low delay, high reward) outnumbered the other 3 options and a poor environment in which the worst option (high delay, low reward) outnumbered the other 3 options. The order of the blocks was counterbalanced between participants.

Participants were exposed to two blocks (“environments,” rich and poor), differing in the frequency of the best (LDHR) and worst (HDLR) options (**Fig. 1c**). In the rich environment, the best option outnumbered the others; in the poor environment, the worst option predominated. Block order was counterbalanced across participants, so that each either completed rich first (“RichPoor”) or the opposite (“PoorRich”). Note that since the two intermediate options, LDLR and HDHR, were identical in profitability (i.e., reward per second) and occurred with identical frequency, to simplify the analysis, we collapse these options into one “intermediate option” category. Separating them does not change the pattern of results (see **Supplementary Material**).

### Participants adapt to global fluctuations in their environment

The MVT predicts that acceptance should depend on an option’s profitability – given our experimental parameters, the best option should always be accepted and the worst option always rejected in both environments – but also on the overall quality of the environment. In particular, it is optimal to accept the intermediate options in the poor environment (where an even worse option predominates), but not in the rich environment (where they just delay frequent encounters with a better alternative).

Accordingly, the decision whether to accept vs. reject an option was sensitive to both its profitability (i.e., reward per second) and also the environment (**Fig. 2a, b**). Specifically, a repeated measures ANOVA on the percentage of accept decisions with option (best, intermediate, worst) and environment (rich, poor) as repeated factors revealed a main effect of environment (Experiment 1: F(1,39) = 15.40, p<0.001; Experiment 2: F(1,37) = 8.38, p<0.01), a main effect of option (Experiment 1: F(2,78) = 540.41, p<0.001; Experiment 2: F(2, 74) = 367.15, p<0.001) and an environment by option interaction (Experiment 1: F(2, 78) = 70.27, p<0.001; Experiment 2: F(2, 74) = 38.11, p<0.001). As predicted by the MVT, in each experiment the interaction was driven by a selective change between environments in acceptance of the intermediate options; this change was greater than the change in acceptance for the best or worst options (intermediate vs. best option, Experiment 1: t(39) = 8.34, p<0.001; Experiment 2: t(37) = 5.60, p<0.001; intermediate vs. worst, Experiment 1: t(39) = 9.89, p<0.001; Experiment 2: t(37) = 7.59, p<0.001, paired sample t-tests on the difference in acceptance rates between environments for intermediate versus either alternative). However, while this effect was in the direction predicted by the MVT (i.e., participants were more selective in the richer environment), the magnitude of the change was not as dramatic as theoretically predicted. This is due in part, as discussed below, to sluggish adjustment in particular circumstances.

**Figure 2.**
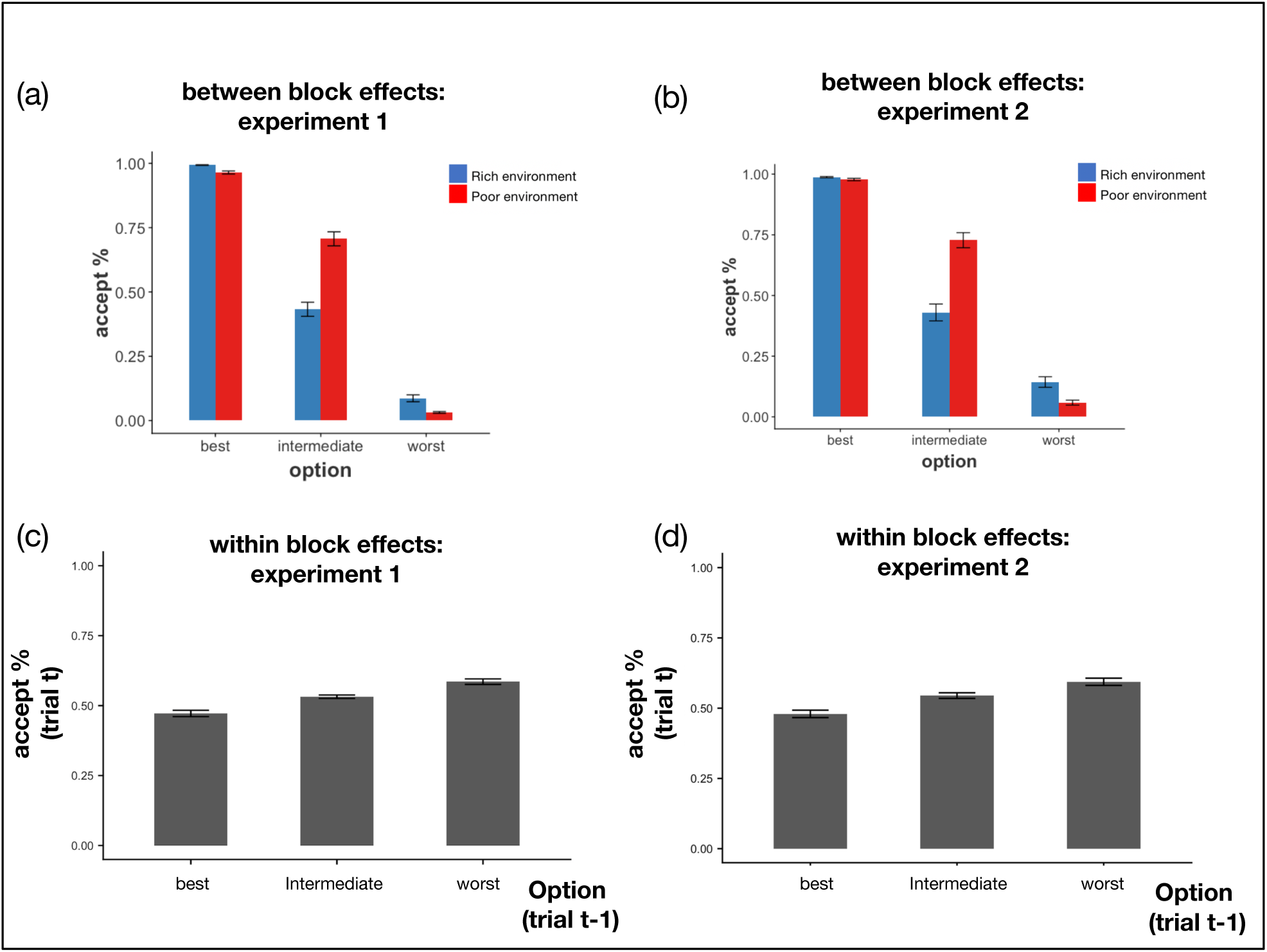
Global and local effects of the environment. **(a), (b)** Participants increased their decision to accept according to how good each option was in terms of it’s reward per second and increased their decision to accept options overall in the poor compared to the rich environment. There was also an environment by option interaction driven by the change in acceptance between environments being greatest for the intermediate options compared to the best (Experiment 1: t(39) = 8.34, p<0.001; Experiment 2: t(37) = 5.60, p<0.001) and worst (Experiment 1: t(39) = 9.89, p<0.001; Experiment 2: t(37) = 7.59, p<0.001, paired sample t-tests on the change in acceptance rates between environments for intermediate versus best/worst options). **(c), (d)** Acceptance rates were modulated by trial to trial dynamics. Participants increased their acceptance rates for an option, the worse the previous option had been (main effect of previous option, controlling for environment: Experiment 1: F(2, 78) = 43.69, p<0.001; Experiment 2: F(2, 74) = 31.68, p<0.001). Best = High Reward, Low Delay option; intermediate = Low Reward, Low Delay and High Reward, High Delay options combined; Worst = Low Reward, High Delay option *Error bars represent standard error of the mean*

### Participants adapt to local fluctuations in their environment

Since the quality of environments was not explicitly instructed, the foregoing results imply that subjects learn from experience how selective to be. Although the MVT itself does not prescribe dynamics, it does immediately suggest a simple class of learning model (Constantino and Daw, 2015; McNamara and Houston, 1985): dynamically estimate the reward rate in the environment (e.g., by an incremental running average), and then use this as an acceptance threshold in the MVT choice rule. Such learning predicts that choices should be sensitive not just to block-wise manipulation of the environmental richness, but also to local variation in offers, since for instance receiving a poor offer will incrementally decrease the estimated reward rate, and incrementally decrease selectivity on the very next trial.

Accordingly, we examined evidence for such trial-to-trial learning by investigating whether the decision to accept an option fluctuated according to recent experience. We separated trials in each environment according to the option participants encountered on the *previous* trial, independent of the decision on the previous trial. To ensure any effect of local context was not merely driven by the block-wise environment type effect discussed above, we controlled for block type by entering environment (rich, poor) as a repeated factor in the analysis along with previous offer (best, intermediate, worst). This analysis revealed that participants increased their acceptance rates (over all options) on the current trial, the worse the option on the previous trial had been (main effect of previous option Experiment 1: F(2,78) = 43.69, p<0.001; Experiment 2: F(2,74) = 31.68, p<0.001, **Fig. 2c**,**d**). In other words, participants became more selective (less likely to accept the current option) immediately following evidence that the environment was rich (previous encounter with the best option) and less selective when it was poor (previous encounter with the worst option), consistent with an MVT-inspired learning model.

### Evidence of block-wise learning

One additional, more global, piece of evidence speaking to the learning process was apparent in block order effects. In particular, we examined whether global fluctuations in acceptance rates were modulated by the order in which the environments were encountered. We did this by implementing a new repeated measures ANOVA with option (best, intermediate, worst) and environment (rich, poor) as repeated factors, and this time included order condition (RichPoor, PoorRich) as a between-participant factor. This revealed an interaction between environment and order condition (Experiment 1: F(1, 38) = 12.11, p<0.01; Experiment 2: F(1, 36) = 7.24, p<0.05) as well as, as before, main effects of environment (Experiment 1: F(1, 38) = 18.22, p<0.01; Experiment 2: F(2, 35) = 8.00, p<0.01) and option (Experiment 1: F(2, 76) = 555.15, p<0.001; Experiment 2: F(2, 72) = 402.48, p<0.001).

The interaction reflected PoorRich participants (those who encountered the poor environment first) being more sensitive to the change in environments, compared to RichPoor participants. That is, PoorRich participants (compared to the opposite order) showed a greater increase in their percentage of accept decisions in the poor environment compared to the rich environment (**Supplementary Fig. 1**). To better visualize this, we calculated a difference score for the change in acceptance rates (across all options, see **Methods**) between the poor environment and rich environment for each participant. Positive scores indicate an overall increase in acceptance rates in the poor environment relative to the rich environment. We then compared these difference scores for participants in the RichPoor condition versus the PoorRich condition (**Fig. 3a**,**b**). This revealed that PoorRich participants showed a significant increase in their percentage of accept decisions in the poor environment compared to the rich environment (Experiment 1: t(20) = 6.79, p<0.001; Experiment 2: t(20) = 4.99, p<0.001, one sample ttest on the difference scores versus 0) in contrast to RichPoor participants who exhibited only a marginally higher rate of acceptance in the poor compared to the rich environment in Experiment 1 (t(18) = 1.97, p=0.07) which was not significant in Experiment 2 (t(16) = 1.17, p=0.26) with there being a significant difference between these two groups in the difference scores (Experiment 1: t(38) = −3.41, p<0.01; Experiment 2: t(36) = −2.08, p<0.05, independent sample ttests, comparing difference scores for RichPoor versus PoorRich participants).

**Figure 3.**
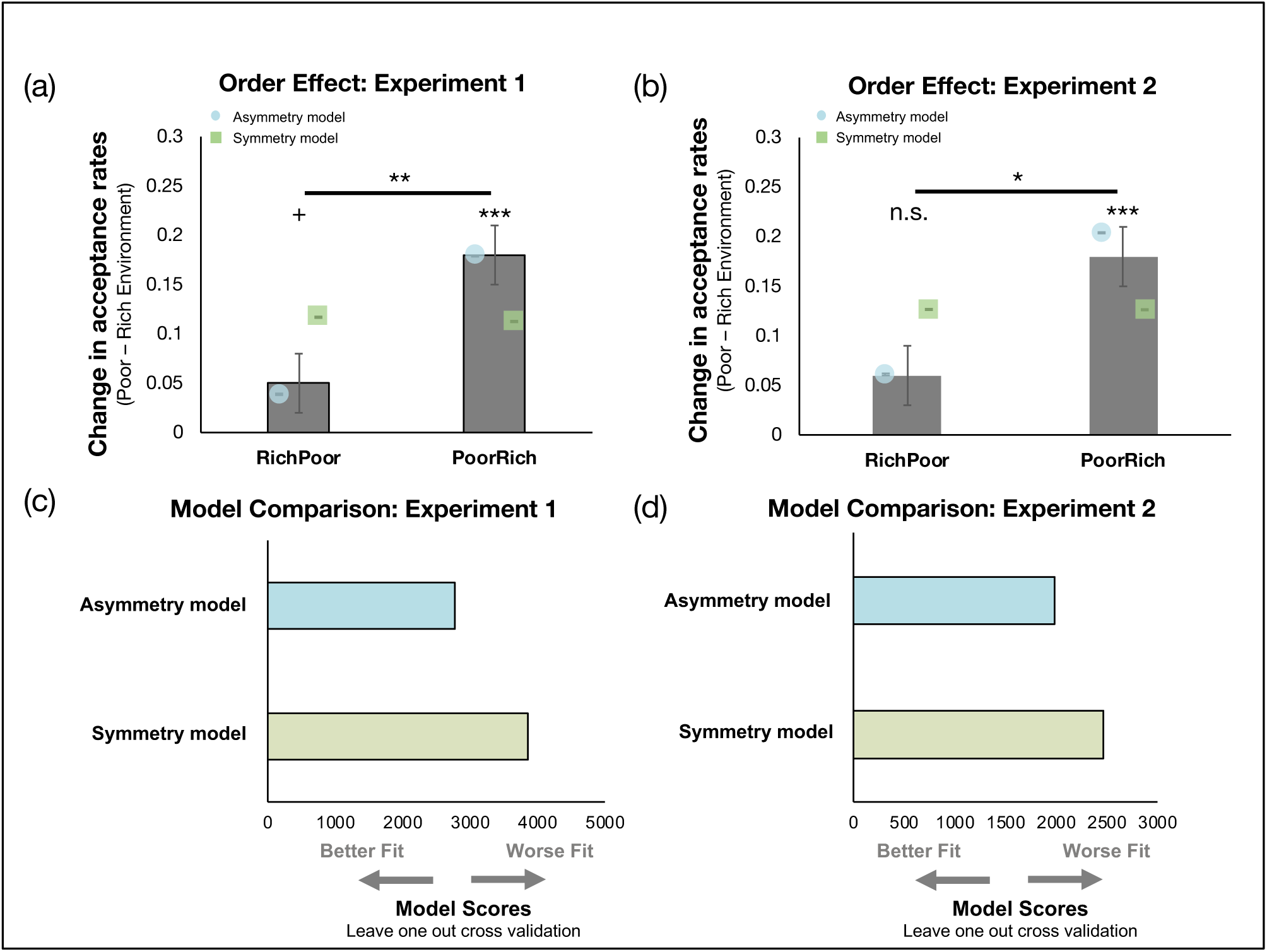
(a) Participants that experienced the poor environment followed by the rich environment (PoorRich group) changed choices between environments to a greater extent than participants that experienced a rich environment followed by a poor environment (t(38) = −3.41, p<0.01, independent ttest comparing the change in acceptance rates between these two groups of participants). The Asymmetry Learning Model was able to recapitulate this order effect by having separate learning rate for when reward rate estimates increased and when they decreased. The Symmetry Model which had a single learning rate predicted that the change in acceptance rates between environment ought to be the same for RichPoor and PoorRich participants. Grey bars indicate participant data. Blue circles represent the pattern of choices generated by simulations from the Asymmetry Model. Green squares represent the pattern of choices generated by simulations from the Symmetry Model. (b) This pattern of results replicated in a second experiment (t(36) = −2.08, p<0.05, independent ttest comparing the change in acceptance rates between these two groups of participants). (c) The Asymmetry Model provided a superior fit to the data than the MVT Model in experiment 1 (t(39) = 7.50, p<0.001, paired sample ttests comparing LOOcv scores for the Asymmetry versus the MVT Model. (d) The Asymmetry Model provided a superior fit to the data than the MVT Model in experiment 2 as well (t(37) = 4.96, p<0.001, paired sample ttests comparing LOOcv scores for the Asymmetry versus the MVT Model). *Error bars represent standard error of the mean* 0.05<p<0.10 *p<0.05; ** p<0.01; *** p<0.001: independent sample ttest / one sample ttest (vs 0) as appropriate *n.s.* = non significant

### Computational modelling

We reasoned that this asymmetry reflected the operation of the underlying learning rule by which subjects adjusted their behavior, trial-by-trial, from the poor to the rich environment or vice versa. For instance, a model that defined an MVT-like threshold from a simple incremental running average of recent events (Constantino and Daw, 2015) would treat either direction of change symmetrically. We hypothesized, however, that the difference could arise from asymmetric learning, as has been studied in other decision domains (Eil and Rao, 2011; Garrett and Sharot, 2014, 2017; Garrett et al., 2014, 2018; Korn et al., 2012; Kuzmanovic and Rigoux, 2017; Kuzmanovic et al., 2015, 2016; Lefebvre et al., 2017): individuals changing their subjective estimates of the environment’s reward rate differentially depending whether the update was in a positive or negative direction. To formally test this, we adapted a computational model (Constantino and Daw, 2015) to capture how participants maintained ongoing estimates of the environments reward rate during the task and tested whether endowing this model with the capacity to shift estimates up and down at different rates depending on whether the information received suggested that the environment was improving (as would be the case when transitioning from poor to rich) versus deteriorating (as would be the case when transitioning from rich to poor) could account for the relative hesitance of participants to change choices when the environment became worse.

Accordingly, we fit the behavioral data in each experiment to two reinforcement learning models (Sutton and Barto, 1998): a *Symmetric Model* and an *Asymmetric Model*. In each of these models, participants were assumed to maintain an ongoing estimate of the reward rate (reward per second), *ρ*, which updated every second. This estimate was then used on each trial to calculate the opportunity cost of accepting an option (*ρ* multiplied by the option’s time delay) at the time of choice. Under this model specification, even though the *absolute cost* (in terms of number of seconds delay) imposed by accepting a specific option was the same throughout the task, the *opportunity cost* could vary according to participant’s current estimate of the environment’s reward rate. The decision to accept or reject the option was modelled as a comparison between the opportunity cost of accepting an option (effectively the value of rejecting) against the reward that the option would collect (effectively the value of accepting). This was implemented using a softmax decision rule with an inverse temperature parameter (β_1_) governing the sensitivity of choices to the difference between these two quantities, and an intercept (β_0_) capturing any fixed, overall bias toward or against acceptance (see **Methods** for further details on the model fitting procedure).

Both models used a delta-rule running average (Rescorla and Wagner, 1972) to update *ρ* according to positive and negative prediction errors. Negative prediction errors were generated every second that elapsed without a reward (for example, each second of a time delay). Positive prediction errors were generated on seconds immediately following a time delay when rewards were received. The difference between the Symmetric Model and the Asymmetric Model was whether there were one or two learning rate parameters. The Symmetric Model contained just a single learning parameter, α. This meant that *ρ* updated at the same rate regardless of whether the update was in a positive or a negative direction. The Asymmetric Model had two learning parameters: α^+^ and α^−^. This enabled *ρ* to update at a different rate, according to whether the update was in a positive (α^+^) or a negative (α^−^) direction. This causes an overall (asymptotic) bias in *ρ* (e.g., if α^+^ > α^−^, then the environmental quality is overestimated), but also dynamic, path-dependent effects due to slower adjustment to one direction of change over the other.

### Asymmetric Model is a better fit to trial-by-trial choice data

We fit the per-participant, per-trial choice timeseries to each model, and compared models by computing unbiased per-participant marginal likelihoods via subject-level Leave One Out cross validation (LOOcv) scores for each participant. The Asymmetric Model provided a superior fit to the choice data than the Symmetric Model both in Experiment 1 (t(39) = 7.50, p<0.001, paired sample ttests comparing LOOcv scores for the Asymmetry versus the Symmetric Model, **Fig. 3c** and **Table 1**) and in Experiment 2 (t(37) = 4.96, p<0.001, **Fig. 3d** and **Table 1**). At the population level, formally comparing the two learning rates (see **Methods**) in the Asymmetric Model revealed that this asymmetry was, on average, significantly biased toward α^+^ > α^−^ (experiment 1: z = 5.50, p<0.001; experiment 2: z = 3.99, p<0.001, **Table 1**). This meant that prediction errors that caused *ρ* to shift upwards (following receipt of a reward) had a greater impact than prediction errors that caused *ρ* to shift downwards (following the absence of a reward). Individually, 90% of participants in experiment 1 and 87% of participants in experiment 2 were estimated to have a higher learning parameter for positive (α^+^) compared to negative (α^−^) errors.

**Table 1:** model fitting and parameters across the three experiments. The table summarizes for each model its fitting performances and its average parameters: LOOCV: leave one out cross validation scores, summed over participants; α: learning rate for both positive and negative prediction errors (Symmetric Model); α+: learning rate for positive prediction errors; α-: average learning rate for negative prediction errors (Asymmetric Model); β_0_: softmax intercept (bias towards reject); β_1_: softmax slope (sensitivity to the difference in the value of rejecting versus the value of accepting an option). Data are expressed as mean and 95% confidence intervals (calculated as the sample mean +/- 1.96* standard error). *\* * * P<0.001 comparing LOOCV scores between the two models, paired sample ttest*

| Experiment / Model | LOOCV | $\alpha$ | $\alpha^+$ | $\alpha^-$ | $\beta_0$ | $\beta_1$ |
| --- | --- | --- | --- | --- | --- | --- |
| **Experiment 1 (N=40)** |  |  |  |  |  |  |
| Symmetric Model | 3857.65 | 0.0027<br>95% CI =<br>[0.007, 0.082] | - | - | 1.03<br>95% CI =<br>[0.71, 1.35] | 0.08<br>95% CI =<br>[0.074, 0.09] |
| Asymmetric Model | 2774.75*** | - | 0.0054<br>95% CI =<br>[0.003, 0.009] | 0.0017<br>95% CI =<br>[0.0009, 0.003] | -1.84<br>95% CI =<br>[-2.58, -1.09] | 0.07<br>95% CI =<br>[0.061, 0.079] |
| **Experiment 2 (N=38)** |  |  |  |  |  |  |
| Symmetric Model | 2467.18 | 0.027<br>95% CI =<br>[0.008, 0.074] | - | - | 0.84<br>95% CI =<br>[0.57, 1.12] | 0.07<br>95% CI =<br>[0.068, 0.080] |
| Asymmetric Model | 1979.60*** | - | 0.0088<br>95% CI =<br>[0.004, 0.017] | 0.0035<br>95% CI =<br>[0.001, 0.010] | -1.21<br>95% CI =<br>[-1.91, -0.52] | 0.07<br>95% CI =<br>[0.067, 0.082] |
| **Experiment 3 (N=38)** |  |  |  |  |  |  |
| Symmetric Model | 3056.73 | 0.12<br>95% CI =<br>[0.05, 0.24] | - | - | 0.77<br>95% CI =<br>[0.49, 1.05] | 0.08<br>95% CI =<br>[0.066, 0.085] |
| Asymmetric Model | 2656.93*** | - | 0.024<br>95% CI =<br>[0.015, 0.035] | 0.012<br>95% CI =<br>[0.008, 0.018] | -1.14<br>95% CI =<br>[-1.65, -0.63] | 0.07<br>95% CI =<br>[0.05, 0.08] |

### Asymmetric Model accounts for order effect

Next, we examined the impact of this asymmetry when the environment changed. Importantly, the model carried *ρ* over between environments rather than resetting it at the start of a new block. Accordingly, simulated participants had to “unlearn” that they were no longer in a poor (PoorRich group) or rich (RichPoor group) environment. Given that the learning asymmetry was revealed to be in a positive direction (α^+^ > α^−^) we reasoned that this unlearning ought to occur faster when going from a poor environment into a rich environment, as large rewards become more commonplace (i.e. the PoorRich group), which would explain the block order effect.

Indeed, simulating the experiment using a population of subjects drawn according to the best-fitting parameter distributions for each model (**Fig. 3a, b**), we found that both models reproduced an overall effect of environment type, but only the Asymmetric Model captured the block order effect. Returning to the fits to actual participants’ choices, we further unpacked the model’s account of this effect by extracting trial by trial estimates of *ρ* from the model’s fit to each trial and participant. The comparison of these estimates between participants in the RichPoor and PoorRich conditions reveals a pattern in accord with the intuition that the global effect arose from slower adjustment of the acceptance threshold by the RichPoor group. In particular, there was a significant environment by condition interaction (Experiment 1: F(1,38) = 13.92, p<0.01, **Fig. 4a**; Experiment 2: F(1, 36) = 18.87, p<0.001, **Supplementary Fig 2a**). This arose out of a significant difference in *ρ* between environments for participants in the PoorRich condition (Experiment 1: t(20) = 8.59, p<0.001; Experiment 2: t(20) = 6.59, p<0.001, paired sample ttest on each participant group separately) which was absent among RichPoor participants (Experiment 1: t(18) = 1.81, p=0.25; Experiment 2: t(16) = 0.98, P=0.34).

**Figure 4.**
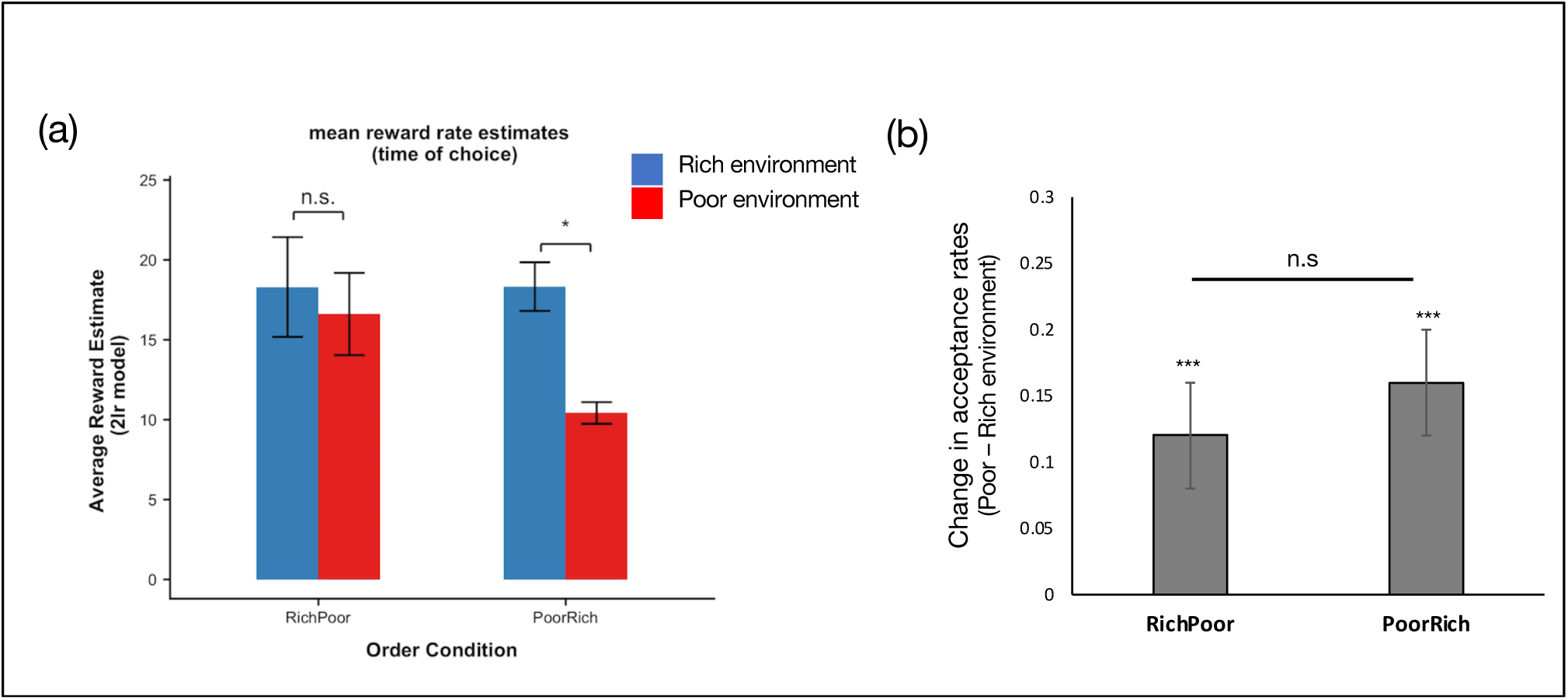
(a) Extracting reward rate estimates (*ρ*) from the Asymmetry model for each participant in each experimental block in Experiment 1 revealed significant environment by condition interaction (F(1, 38) = 13.92, p<0.01). This arose out of a significant difference in *ρ* between environments for participants in the PoorRich condition (t(20) = 8.59, p<0.001, paired sample ttest) which was absent among participants assigned to the RichPoor condition (t(18) = 1.81, p=0.25). (b) In Experiment 3, both groups of participants significantly changed their acceptance rates between environments (RichPoor: t(14) = 3.07, p<0.01; PoorRich: t(22) = 4.46, p<0.001) and there was no longer any difference in the change in acceptance rates between RichPoor and PoorRich participants (t(36)=−0.77, p = 0.45, independent sample ttest). * p<0.05, paired sample ttest; n.s. = non significant (p>0.05); * * * p<0.001, one sample ttest (vs 0)

### Order effect disappears given sufficient exposure to a poor environment

To further probe whether asymmetric learning caused strategies to perseverate among RichPoor participants, we ran a third cohort of participants. The procedure was exactly as for Experiments 1 and 2 but with one key difference. Now, between the two environments (either between the Rich and the Poor environment or between the Poor and the Rich environment) we sandwiched a third environment in which participants only saw the worst option (HDLR, see **Methods** for full details). We predicted that with the addition of this environment, participants in the RichPoor condition would now have sufficient exposure to a lean environment to allow their expectations to adjust before entering the poor environment. In doing so, we predicted that we ought no longer to observe a difference in the change in acceptance rates between environments for RichPoor and PoorRich participants.

The Asymmetric Model once again proved a better fit to choice behavior compared to Symmetric Model (t(37) = 5.64, p<0.001, paired sample ttests comparing LOOcv scores for the Asymmetric Model versus the Symmetric Model, see also **Table 1**) with α^+^ being greater than α^−^ (z = 4.61, p<0.001). But in contrast to before, both groups significantly changed their acceptance rates between environments (RichPoor: t(14) = 3.07, p<0.01; PoorRich: t(22) = 4.46, p<0.001) and there was no difference in this change in acceptance rates between RichPoor and PoorRich participants (t(36)=−0.77, p = 0.45, independent sample ttest, **Fig. 4b**).

### Preference between intermediate options

The two intermediate options, LDLR and HDHR, were equated in terms of the reward they provided per second and were encountered with equal frequency in each environment (**Fig. 1c**). Nevertheless, in experiments 1 and 2, participants did not show an equal preference for these two options, generally preferring LDLR to HDHR (Experiment 1: Mean Difference in Acceptance rates = 24%, t(39) = 3.95; Experiment 2: Difference in Acceptance rates = 12%, t(37) = 2.20, p<0.05, paired sample ttest on acceptance rates for LDLR versus HDHR averaged across both environments. See also **Supplementary Figure 1**). This preference did not replicate significantly in the third experiment but the pattern was in the same direction (Experiment 3: t(37) = 1.27, p>0.05). This pattern of results was also recovered in simulations from the Asymmetric Model (Experiment 1: Difference in Acceptance rates = 22%, t(999) = 125.35, p<0.001; Experiment 2: Difference in Acceptance rates = 15%; t(999) = 93.87, p<0.001, paired sample ttest comparing LDLR versus HDHR acceptance rates, averaged over both environments from the Asymmetric Model simulations).

## Discussion

Across three experiments using a prey selection task, we find that individuals adjusted their choices both between environments – becoming less selective in a poor compared to a rich environment – and within environments – altering selectivity on a trial by trial basis according to the previous encounter. The block-wise effect is consistent with foraging theory (Stephens & Krebs, 1986) and previous empirical data both from animals (Charnov, 1976; Freidin & Kacelnik, 2011; Hayden et al., 2011; Kacelnik, 1984; McNickle & Cahill, 2009; Stephens & Krebs, 1986) and humans (Hutchinson et al., 2008; Jacobs & Hackenberg, 1996; Kolling et al., 2012; McCall, 1970; Smith & Winterhalder, 1992). The trial-wise effect is a rather direct prediction of incremental learning rules for the acceptance threshold, which have previously been proposed and studied in terms of a different class of foraging tasks, patch leaving tasks (Constantino and Daw, 2015; McNamara and Houston, 1985; Zhang et al., 2015), where choice-by-choice effects are harder to observe due to the task structure.

However, inconsistent with optimal foraging theory, we observed interesting suboptimalities in individuals’ choices. In particular, adjustments in choices between environments were attenuated when an individual’s environment became worse compared to when the environment improved. The experience-dependent nature of this bias strongly suggests that it is rooted in learning, and we present modeling showing that it can be understood in terms of an asymmetric learning rule, which scales positive and negative prediction errors differently. Such decomposition of learning by valence is a recurring theme in decision neuroscience, but its consequences in the foraging setting – path-dependent biases in estimates of opportunity cost, leading to systematic choice biases – have not previously been appreciated. Indeed, optimism biases of the sort implied by these models may have particularly important effects in many real-world foraging-like tasks (including hiring and employment decisions and mate selection) because these learned estimates play such a key role in choice: encountering opportunities serially forces the decision-maker to compare them to a learned (and potentially biased) estimate of the “other fish in the sea” rather than directly to their alternatives.

We were able to account for the deviations from foraging theory by augmenting a learning rule that had previously been used in the patch foraging setting (McNamara and Houston, 1985; Constantino & Daw, 2015). Across all three experiments, endowing the model with separate learning rates for positive and negative adjustments to the environment’s reward rate provided a better qualitative and quantitative fit to participants’ choices compared to a baseline model which did not distinguish between these two types of adjustments. This feature provided the model with the capacity to shift estimates of the environments reward rate up and down at different rates, enabling it to recapitulate the differences in choice adjustments depending on the sequence in which participants experienced rich and poor environments.

Interestingly, the order effect, and the learning account of it, indicate that participants carry over information about the reward rate from one environment into the next. This is despite the fact that participants are explicitly told at the start of the experiment that they will experience different environments in the task and, during the experiment, new environments are clearly signaled. This stands in contrast to another recent study (Palminteri et al., 2015) in which participants encountered a set of distinct contexts interleaved, and were able to associate distinct reward rate estimates with each context. It may be that also in the task we use here, the introduction of more transitions, more environments, or more volatility, would prompt a shift towards this type of discrimination and mitigate the carryover biases we observe.

Our learning-based account is further supported, at least to a limited extent, by our third experiment, in which the block order effect was not significant in a version of the task that included additional experience with a poor reward rate. This negative finding is predicted by the learning model, and tends to mitigate against other explanations (e.g. ones in which the block order asymmetry relates to some sort of primacy or anchoring on the initial experience) that would predict equivalent effects to the (similarly designed and powered) Experiments 1 and 2. Of course, null effects (even predicted ones) should be interpreted with caution, and the difference in significance between experiments does not necessarily imply the effects are themselves different. Future experiments including both conditions in a single design will be required to permit their direct statistical comparison.

Another, subtler feature of our results is also consistent with the model. Although the core idea of reward rate maximization and the MVT seem to predict equivalence between two options with the same reward rate, participants tended to accept one of our two intermediate options (small rewards quickly) more than another (larger rewards slowly), despite these having equal reward rate. In fact, this asymmetry is also predicted by the model and constrains the form of its choice rule. In particular, the choice rule is expressed in terms of reward amount (the difference between the reward on offer and the opportunity cost, in units of reward, for occupying that time) rather than the alternative of comparing these quantities expressed as rates (normalized by delay). Softmax choice on the former basis results in a larger decision variable and more deterministic choices for the longer-delay option; if the net decision variable is negative (typically the case in our regime when α^+^ > α^−^, because the opportunity cost is overestimated), the shorter one will be rejected less often. Other features not included in the model, such as time discounting, might also contribute to this preference.

These results may reflect a much broader feature of learning. In a variety of other domains, information integration has been revealed to be an asymmetric process in which beliefs are more readily updated in response to information that calls for an adjustment in a positive rather than negative direction (Sharot & Garrett, 2016). For example, people are more likely to update beliefs when receiving ‘good news’ regarding their environment (such as learning their likelihood of being a victim of credit card fraud is lower than they thought) than when receiving ‘bad news’ (learning that their likelihood is greater than they thought) (Garrett and Sharot, 2017; Garrett et al., 2014; Kuzmanovic and Rigoux, 2017; Kuzmanovic et al., 2015, 2016; Moutsiana et al., 2013; Sharot et al., 2011). The same pattern emerges when people receive desirable and undesirable information about their financial prospects (Wiswall and Zafar, 2015), feedback about their intellectual abilities (Eil and Rao, 2011; Mobius et al., 2011), personality (Korn et al., 2012) and physical traits such as attractiveness (Eil and Rao, 2011). In all of these cases, desirable information is integrated into prior beliefs more readily than undesirable information, resulting in positively biased beliefs (Moutsiana et al., 2013; Sharot et al., 2011). Consistent with the results that we observe herein, this phenomenon is thought to arise out of an asymmetry in how individuals weight discrepancies between existing beliefs and new information that calls for beliefs to be revised, with a greater weight being placed on discrepancies that motivate a shift to a more positive set of beliefs about the world compared to a more negative set of beliefs (Garrett and Sharot, 2017; Garrett et al., 2014; Kuzmanovic and Rigoux, 2017; Kuzmanovic et al., 2015, 2016; Moutsiana et al., 2013; Sharot et al., 2011).

Neural accounts of learning have also stressed the separation of negative and positive information and updates, which (given that firing rates cannot be negative) are potentially represented in opponent systems or pathways (Daw et al., 2004; Frank et al., 2004; Collins and Frank, 2014). Appetitive and aversive expectancies, and approach and avoidance, are also (at least on some accounts) associated with engagement of partly or altogether distinct brain regions (Bartra et al., 2013; Bernardi and Salzman, 2017; Garrison et al., 2013; Hayes et al., 2014; Namburi et al., 2016; Palminteri and Pessiglione, 2017; Palminteri et al., 2015). More particularly, the direct and indirect pathways through the basal ganglia have been argued to support distinct pathways for action selection versus avoidance (Frank et al., 2004; Collins and Frank, 2014), with positive and negative errors driving updates toward either channel. These types of models also typically incorporate asymmetric updating, though not necessarily consistently biased in all circumstances toward positive information (α^+^ > α^−^).

One important interpretational caveat, at least with respect to the apparent analogy with other work on biased updating is that, compared to these other cases, in the foraging scenario we use here, upward and downward adjustments in reward rate estimates differ in ways other than valence. Specifically, upward errors are driven by reward receipt and downward ones by the passage of time, events which may be perceived and processed differently in ways that may also contribute to asymmetric learning. Future experiments will be required to uncover if valence – whether a piece of information is good or bad – is the key attribute that gives rise to the asymmetry we observe.

Another question that awaits future work is whether the learning-based mechanism we describe also contributes to biases that have been observed in other foraging scenarios. Notably, a number of other reported biases, especially in patch foraging tasks, tend to reflect over-acceptance or overstaying (Constantino and Daw, 2015, Lenow et al., 2017, Wiekenheiser et al, 2013), which would imply pessimistic rather than the optimistically biased opportunity costs we observe here, and a bias toward negative updating. This, in turn, may relate to a further experimental and theoretical question: what features of experience or environment determine the balance in sensitivity to positive and negative prediction errors (Palminteri et al., 2015)?

The possibility of biased reward rate evaluations, and particularly that they might be pessimistic in some circumstances, may also be of clinical relevance. (Huys et al., 2015) propose that a pessimistic estimate of an environment’s rate of reward may offer a common explanation for a range of diverse symptoms of Major Depressive Disorder such as: anergia, excessive sleeping, lack of appetite and psychomotor retardation. Biased updating of the sort we discuss here might protect against the development of these maladaptive behaviors. Interestingly, the positive learning asymmetries that have been observed previously (in non-foraging related tasks), have been shown to disappear in clinically depressed individuals (Garrett et al., 2014; Korn et al., 2014). Whether the learning asymmetry we observe also disappears or reverses toward a negative direction in MDD patients is an important question for future research.

## Methods

### Participants

A total of 62 participants were recruited for Experiment 1. 22 of these were excluded leaving a final sample of 40 participants (mean ± standard deviation (s.d.) age: 32.73 (7.94); 11 female). A total of 55 participants were recruited for experiment 2. 17 of these were excluded leaving a final sample of 38 participants (mean ± standard deviation (s.d.) age: 33.71 (8.83); 16 female; 2 gender undisclosed). Participants were recruited online via <u>Amazon Mechanical Turk</u>. Following best practice for studies with this population (Crump et al., 2013), several a priori exclusion criteria were applied to ensure data quality.

Participants were excluded if any of the following applied: (1) Did not finish the task (n=5); (2) Made 20 or more missed responses (n=14); (3) Made 10 or more incorrect force trial responses (either accepting options when forced to reject or rejecting options when forced to accept, n=1); (4) Poor discriminability, defined as choosing the worst option (low reward, high delay) a greater percentage of times than the best option (high reward, low delay) or accepting/rejecting all options on every single trial (n=2). Participants were paid $1.5 plus a bonus payment between $1.5 and $4 depending on performance in the task. The study design was approved by Princeton University’s ethics committee.

### Behavioral Task (Fig. 1)

The task began by asking participants to provide basic demographic information (age and gender). After providing this information, participants read task instructions on screen at their own pace and undertook a practice session for four minutes. The two options and the environment used in the training session were not used in the actual task. After completing the training session, participants were then required to attempt a multiple-choice quiz to check their understanding of the task and instructions. Participants needed to get all the questions correct to pass the quiz and continue to the task. Participants who failed the quiz were required to reread the instructions and attempt the quiz again. (They were not asked to retake the training again.)

The main experiment comprised two different environments, each lasting 15 (Experiment 1) or 10 (Experiment 2) minutes in a block design (i.e., participants completed 15/10 minutes of one environment and then switched to the other environment for a new period of 15/10 minutes). Participants were told in the instructions that they would encounter the same options in each environment but the frequency of the options could vary between the environments and that their goal was to gain as collect as much fuel as possible in the time available. There was a break in-between the two environments. The background color was different for each environment and before continuing to the 2^nd^ environment, participants were explicitly told that they would now experience a new environment.

On each trial, one of four stimuli (options) appeared and moved toward one of two targets; one on the left side of the screen and another on the right. Participants accepted an option by selecting the target the option approached or rejected an option by selecting the alternate target. Participants had two seconds to respond by selecting the left/right target using separate keys on their computer keyboard. Following acceptance of an option, participants faced a time delay (1 or 7 seconds depending on the stimulus) during which they were required to keep the key used to accept the option pressed down. During this time the option on screen gradually approached them. At the end of the time delay, the target turned yellow and the number of points obtained appeared above the target (1 second) at which point participants could release the key press. Following rejection of an option, the experiment progressed to the next trial. If the participant failed to respond during the encounter screen or released a keypress before a time delay had finished, they faced a timeout of 8 seconds. This meant that having made the decision to accept an option, it was disadvantageous to then abort.

As an attention check and to encourage learning, 25% of trials were forced choice trials. On these trials, participants saw a red asterisk (*) appear over one of the two targets and had to choose that target. This meant that if it appeared over a target that a stimulus was approaching, they had to accept the stimulus on that trial. If it appeared over a target the stimulus was not approaching, they had to choose to reject the stimulus on that trial. Participants were told that more than 5 incorrect forced choice trials would see their bonus payment reduced by half. At the end of the experiment, participants were told how many points they had amassed in total and their corresponding bonus payment.

The task was programmed in JavaScript using the toolbox jsPsych (Leeuw, 2015) version 5.0.3.

### Stimuli

Four different stimuli provided participants with one of two levels of points (presented to participants as 20%/80% the length of a horizontal bar displayed on the reward screen, **Fig 1a**, which corresponded to 20/80 points) and incurred one of two time delays (2 or 8 seconds including the 1 second for reward screen display). These options therefore assumed a natural ordering in terms of their value from “best” (low delay high reward = LDHR), intermediate (low delay low reward = LDLR, high delay high reward = HDHR) to “worst” (high delay low reward = HDLR). The frequency of each option varied between the two different environments. In the “rich” environment, the best option was encountered 4 times more than the other 3 options. In a sequence of 7 trials, the participant would encounter (in a random order) LDHR 4 times, and also encountering each of LDLR, HDHR and HDLR once. In the “poor” environment, in a sequence of 7 trials, the participant would encounter HDLR 4 times and the other 3 options (LDHR, LDLR and HDHR) once (see **Fig. 1c**).

### Behavioral Analysis

To examine changes in acceptance rates between environments, we calculated the percentage of accept decisions for each participant for each option they saw in each environment. Forced choice trials and missed responses were excluded from this analysis. They are not, however, excluded from the computational models. To simplify the analysis presented in the main paper we collapsed the two intermediate options - which have an identical profitability (i.e. reward per second) of 10 points per second - together. We then entered these percentages into a repeated measures ANOVA with option (best/intermediate/worst) and environment (rich/poor) as repeated factors. We also ran the same ANOVA treating the two intermediate options as separate levels. In this instance, option (LDHR/LDLR/HDHR/HDLR) and environment (rich/poor) were entered as repeated factors. These results are presented in the **Supplementary Material**.

To examine how changes in acceptance rates related to the order with which participants encountered the environment, we entered the percentage of accept decisions into a separate repeated measures ANOVA with option (LDHR/intermediate/HDLR) and environment (rich/poor) as repeated factors. Condition (RichPoor or PoorRich), which indicates which ordering of environments participants faced, was entered as a between participant factor. As above, we also ran the same ANOVA treating the two intermediate options as separate levels. These results are presented in the **Supplementary Material**.

To better characterize order effects, we calculated the difference in acceptance rates between environments (poor minus rich) for each option (LDHR, LDLR, HDHR and HDLR). Hence positive scores indicate an increase in acceptance rates in the poor environment compared to the rich environment. We then calculated the mean change across the 4 options as a measure of overall change in acceptance rate between environments. We then compared the overall change in acceptance scores between participants (RichPoor and PoorRich) using independent sample ttests.

Finally, to examine trial to trial fluctuations, we first partitioned trials according to the option presented (best, intermediate, worst). We then calculated separately for each participant and for each environment, the percentage of times on the *next* trial the decision was to accept. We entered these acceptance scores into a 3 by 2 way repeated measure ANOVA with option (best, intermediate, worst) and environment (rich, poor) as factors. To visualize these fluctuations (**Fig 2c**,**d**), we calculated the average of the two acceptance scores - one score for each environment - for each option.

### Computational Models

The optimal policy from the MVT is to accept an option, *i*, whenever the reward, *r*_*i*_, that the option obtains exceeds the opportunity cost, *c*_*i*_, of the time taken to pursue the option. This opportunity cost (*c_i_*) is calculated as the time, *t*_*i*_, that the option takes to pursue (in seconds) multiplied by the estimated reward rate (per second) of the environment, *ρ*.

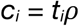

Participants should therefore accept an option whenever:

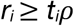

Note we assumed that the quantities *r*_*i*_ and *t*_*i*_ were known to participants from the outset since they were easily observable and each of the 4 options (*i* = {1,2,3,4}) always provided the exact same *r*_*i*_, and *t*_*i*_. But models that dropped this assumption (and instead assumed *r*_*i*_ and t_*i*_ were learned via experience rather than known) provided similar patterns of results (see **Supplementary Material**).

We assumed that subjects learned *ρ* in units of reward, using a Rescorla-Wagner learning rule (Rescorla and Wagner, 1972; Sutton and Barto, 1998) which is applied at every second. After each second, the value of the environment is updated according to the following rule:

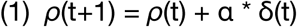

Here *t* indexes time in seconds. δ(t) is a prediction error, calculated as:

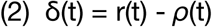

r(t) is the reward obtained. This will either be 0 (for every second in which no reward is obtained, i.e. during search time, handling time and timeouts from missed responses) or equal to *r*_*i*_ (following receipt of a reward).

The learning rate *α* acts as a scaling parameter and governs how much participants change their estimate of the reward rate of the environment (*ρ*) from one second to the next. This estimate increases when *r* is positive (i.e. when a reward is obtained) and decreases every second that elapses without a reward.

We implemented two versions of this reinforcement learning model. A *Symmetric Model*, with only a single *α* and a modified version, an *Asymmetric Model*, which had two *α*: *α*^*+*^ *and α*^−^. In this second model, updates to *ρ* apply as follows:

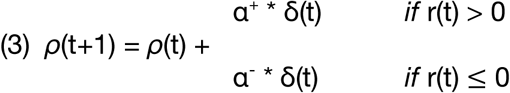

This second model allows updates to occur differently according to whether a reward is received in the environment or not. This is close to identical to updating *ρ* contingent on whether the prediction error, δ(t), is positive or negative (Frank et al., 2007; Lefebvre et al., 2017; Niv et al., 2015), as for the vast majority of trials (95%) in which r(t) > 0, it was also the case that δ(t) > 0 whilst in all trials (100%) in which r(t) ≤ 0, it was also the case that δ(t) < 0. We refer to the mean difference in learning rates as the *learning bias* (α^+^ – α^−^). A positive learning bias (α^+^ > α^−^) indicates that participants adjust their estimates of the environments reward rate to a greater extent when a reward is obtained compared to when rewards are absent. The converse is true when the learning bias is negative (α^+^ < α^−^). If there is no learning bias (α^+^ = α^−^) then this model is equivalent to the simpler Symmetric Model with a single α.

To account for both the delay and the reward received in the final second of handling time (when participants received a reward), this was modelled as two separate updates to *ρ*; one update from the delay (in which δ(t) = 0 – *ρ*(t)) followed by a second update from the reward received (in which δ(t) = r_i_ – *ρ*(t)). Swapping the order of these updates or omitting the first update (from the delay) altogether, did not alter the pattern of results.

The probability of choosing to accept an option is estimated using a softmax choice rule (implemented at the final (2^nd^) second of the encounter screen as follows:

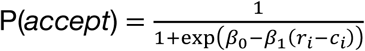

This formulation frames the decision to accept an option as a stochastic decision rule in which participants (nosily) choose between two actions (accept/reject) according to the respective value of each of them. The temperature parameter β_**1**_ governs participants’ sensitivity to the difference between these two values whilst the bias term β_**0**_ captures a participant’s general tendency towards accepting/rejecting options (independent of the values of each action). Note that under this formulation, negative values for β_**0**_ indicate a bias towards accepting options, positive values indicate a bias towards rejecting options.

At the beginning of the experiment, *ρ* was initialized to the average (arithmetic) reward rate across the experiment (Constantino and Daw, 2015). (Although clearly not realistic as a process level model, this was included for simplicity to avoid estimation pathologies and special-case model features associated with initial conditions.) In subsequent environments, the average reward rate carried over from the previous environment. In other words, there was no “resetting” when participants entered a new environment. Rather, they had to unlearn what they had learnt in the previous environment.

For each participant, we estimated the free parameters of the model by maximizing the likelihood of their sequence of choices, jointly with group-level distributions over the entire population using an Expectation Maximization (EM) procedure (Huys et al., 2011) implemented in the Julia language (Bezanson et al., 2012), version 0.7.0. Models were compared by first computing unbiased per subject marginal likelihoods via subject-level cross validation and then comparing these likelihoods between models (Asymmetric versus Symmetric) using paired sample ttests.

To formally test for differences in learning rates (α^+^, α^−^) we estimated the covariance matrix 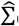 over the group level parameters using the Hessian of the model likelihood (Oakes, 1999) and then used a contrast 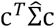 to compute the standard error on the difference α^+^ – α^−^.

### Simulations

To examine the qualitative fit of each model to the data we ran simulations for both the Symmetric Model and the Asymmetric Model. For each simulation (n=1000), we ran a group of 40 virtual participants (n=40). For each virtual participant, we assigned a set of parameters (β_0_, β_1_ and α in the case of the Symmetric Model; β_0_, β_1_, α^+^ and α^−^ in the case of the Asymmetric Model) by randomly selecting from the best fit parameters generated by the computational model (fit to actual participants choices) and randomly assigned the order with which they encountered the environments (either rich to poor *or* poor to rich). Then, we simulated the learning process by which *ρ* evolved as options were sequentially encountered and stochastically accepted/rejected. The learning process was exactly as described for the respective computational models. Crucially, *ρ* was initialized exactly as for the model and allowed to carry over between blocks. We then calculated the acceptance rates for each virtual participant for each option in each block of trials. We then calculated the average acceptance rates over all participants in the simulation as well as separately for participants assigned to each order condition. We then averaged these 3 sets of acceptance rates over each of the simulations.

### Participants and Procedure Experiment 3

A total of 59 participants were recruited for Experiment 3. 21 of these were excluded (identical exclusion criteria as for Experiments 1 and 2) leaving a final sample of 38 participants (mean ± standard deviation (s.d.) age: 34.63 (8.58); 12 female). The experiment was exactly as described as for Experiment 1 and Experiment 2 with one key difference. In this version there were three environments instead of two. A rich environment and a poor environment (exactly as for Experiment 1 and 2) as well as a 3^rd^ “HDLR” environment. The only options participants encountered in the HDLR environment were the worst (HDLR) option. Participants had 10 minutes in each environment. The ordering was either Rich, HDLR, Poor or Poor, HDLR, Rich.

## Data Availability

All data and code are available on request to the authors.

## Acknowledgements

We would like to thank Scott Grafton and Neil Dundon for helpful insight and discussions.

## Supplementary Material

### Additional Behavioral Analysis

#### Block wise learning and order effects

In the main paper, to simplify the analysis, we collapsed the two intermediate options (LDLR and HDHR) into one intermediate category. The same pattern of results is found however if these are treated separately along with the other two options (LDHR and HDLR) (**Supplementary Fig. 1**). Specifically, a repeated measures ANOVA looking at the percentage of accept decisions with option (LDHR, LDLR, HDHR, HDLR) and environment (rich, poor) as repeated factors revealed a main effect of environment (Experiment 1: F(1,39) = 31.75, p<0.001; Experiment 2: F(1,37) = 16.44, p<0.001; Experiment 3: F(1, 37) = 29.63, p<0.001), a main effect of option (Experiment 1: F(3,117) = 154.17, p<0.001; Experiment 2: F(3, 111) = 133.65, p<0.001; Experiment 3: F(3, 111) = 117.82, p<0.001) and an environment by option interaction (Experiment 1: F(3,117) = 26.47, p<0.001; Experiment 2: F(3,35) = 25.25, p<0.001; Experiment 3: F(3,35) = 32.24, p<0.001).

To separately examine differences in blockwise learning between participants we ran the same ANOVA with option (LDHR, LDLR, HDHR, HDLR) and environment (rich, poor) as repeated factors but now also included order condition (RichPoor, PoorRich) as a between subjects’ factor. As reported in the main text (where the intermediate options are collapsed), this revealed an interaction between environment and order condition in Experiment 1 (F(1,38) = 11.64, p<0.01) and Experiment 2 (F(1,36) = 4.33, p<0.05). As expected, this interaction was not significant in Experiment 3 (F(1,36) = 0.57, p=0.45).

### Additional Computational Models

#### Learn Options Models

The Symmetric Model and Asymmetric Model described in the main text assume that the rewards and time investment (*r*_*i*_ and *t*_*i*_) of each of the 4 options (*i* = {1,2,3,4}) were known from the outset. This seems a plausible assumption since the options were visually very distinct, outcomes (rewards and delays) easily observable and were stationary (i.e. each option always provided the exact same *r*_*i*_, and *t*_*i*_). Nonetheless, it may be the case that individuals learnt the rewards and time investments associated with each option over time following feedback. To test whether this was the case, we augmented the Symmetric Model and Asymmetric Model so that the reward and time investment of each options was learnt. Specifically, we initialized a set of reward (Qr_i_) and time (Qt_i_) Q values for each option to 0 (where i indexes the option from 1 to 4). Each of these Q values was then updated following acceptance of an option according to two delta rules (one for Qr and one for Qt) as follows:

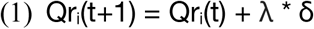

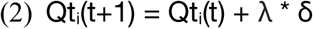

δ is a prediction error, calculated for reward and time respectively as:

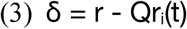

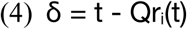

where r is the reward received and t is the time investment required following acceptance of an option. λ is the learning rate used to update estimates of the reward and time investment associated with each option (we use λ rather than α to distinguish it from the learing rate used to update estimates of the environments reward rate).

The opportunity cost (*c*_*i*_), rather than being the product of the actual time investment required and the estimated reward rate (per second) of the environment (*ρ*) is now calculated as the current estimate of the time that the option takes to pursue (Qt_i_) multiplied by the estimated reward rate (per second) of the environment (*ρ*):

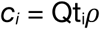

The decision to accept or reject, rather than being modelled as the difference between the estimated opportunity cost (c_i_) and the actual reward an option will gain if accepted (r), is now calculated as the difference between the estimated opportunity cost and the current estimate of the reward that the option will gain (Q_r_). As before, this was implemented in a softmax function:

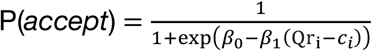

We implemented this extra learning component for both the Symmetric and the Asymmetric Model (for each experiment) with everything else exactly as described for these models. In each case we used a different learning rate (λ) to model learning of the rewards/costs associated with each option to the learning rate(s) used to model learning of the environments rate of reward (α in the case of the Symmetric Model, α^+^ and α^−^ in the case of the Asymmetric Model).

In each experiment, as we found previously (when *r_i_* and *t_i_* were assumed to be correctly known from the outset rather than learnt following feedback), a model with two learning rates for the environments rate of reward provided a better fit to the data compared to a model where this rate was updated using a single learning rate (Symmetric Learn Options Model). The asymmetry between α^**+**^ and α^−^ was again biased in a positive direction in each experiment (Experiment 1: z = 5.90, p<0.001; Experiment 2: z = 5.67, Experiment 3: z = 6.11, p<0.001, **Supplementary Table 1**).

**Supplementary Table 1:**
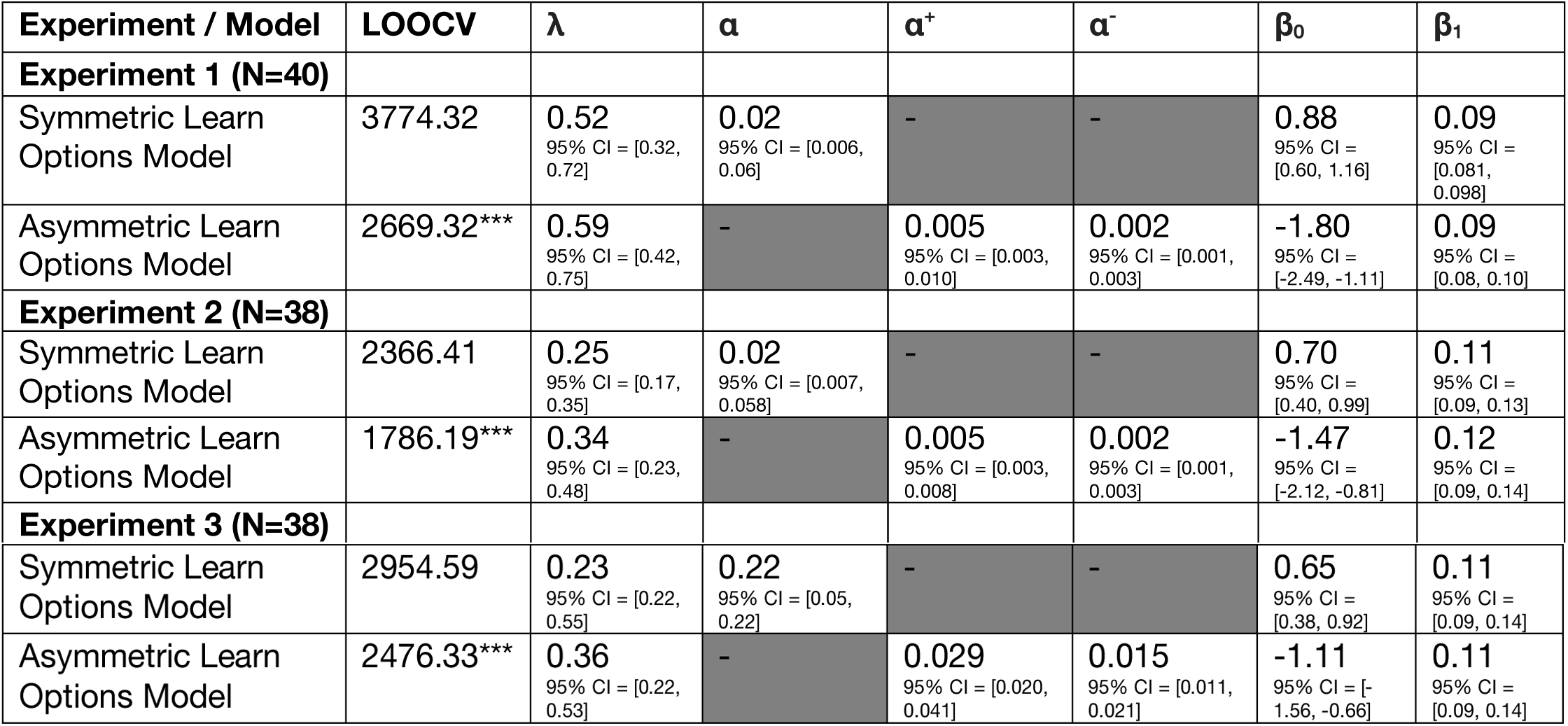
model fitting and parameters for the Learn Option models for each experiment. model fitting and parameters across the three experiments for models which incorporate learning of the rewards and time investment associated with each option. The table summarizes for each model its fitting performances and its average parameters: LOOCV: leave one out cross validation scores, summed over participants; α: learning rate for both positive and negative prediction errors (Symmetric Learn Options Model); α+: learning rate for positive prediction errors; α-: average learning rate for negative prediction errors (Asymmetric Learn Options Model); λ: learning rate for rewards and time investment associated with each option; β_0_: softmax intercept (bias towards reject); β_1_: softmax slope (sensitivity to the difference in the value of rejecting versus the value of accepting an option). Data are expressed as mean and 95% confidence intervals (calculated as the sample mean +/- 1.96* standard error). *\* * * P<0.001 comparing LOOcv scores between each model and the Asymmetric Model, paired sample ttest*

**Supplementary Figure 1.**
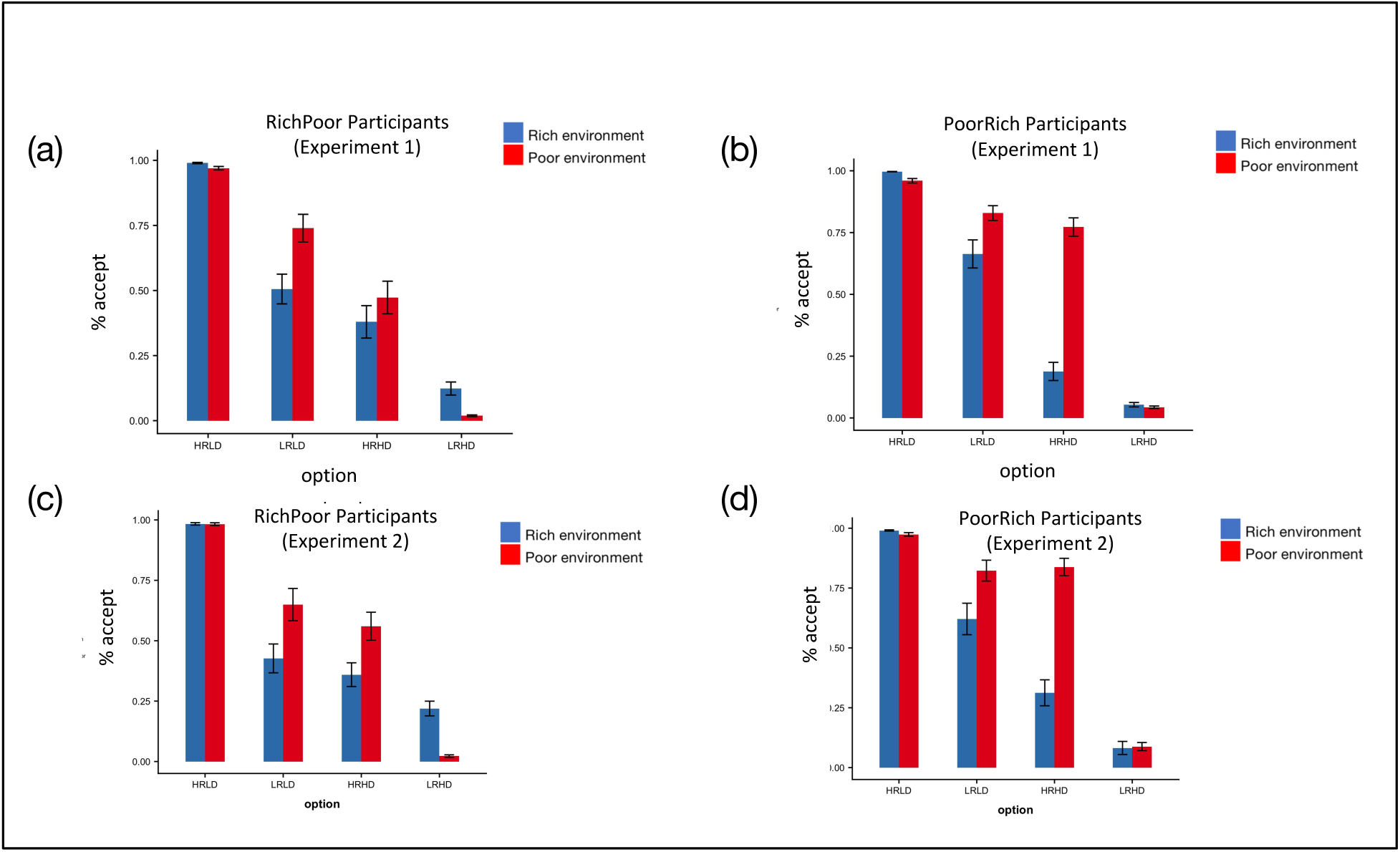
Acceptance rates (%) for each option in each environment separately for each group of participants (RichPoor, PoorRich). As reported in the main text, there was an interaction between environment and order condition in Experiment 1 (F(1,38) = 11.64, p<0.01) and Experiment 2 (F(1,36) = 4.33, p<0.05).

**Supplementary Figure 2.**
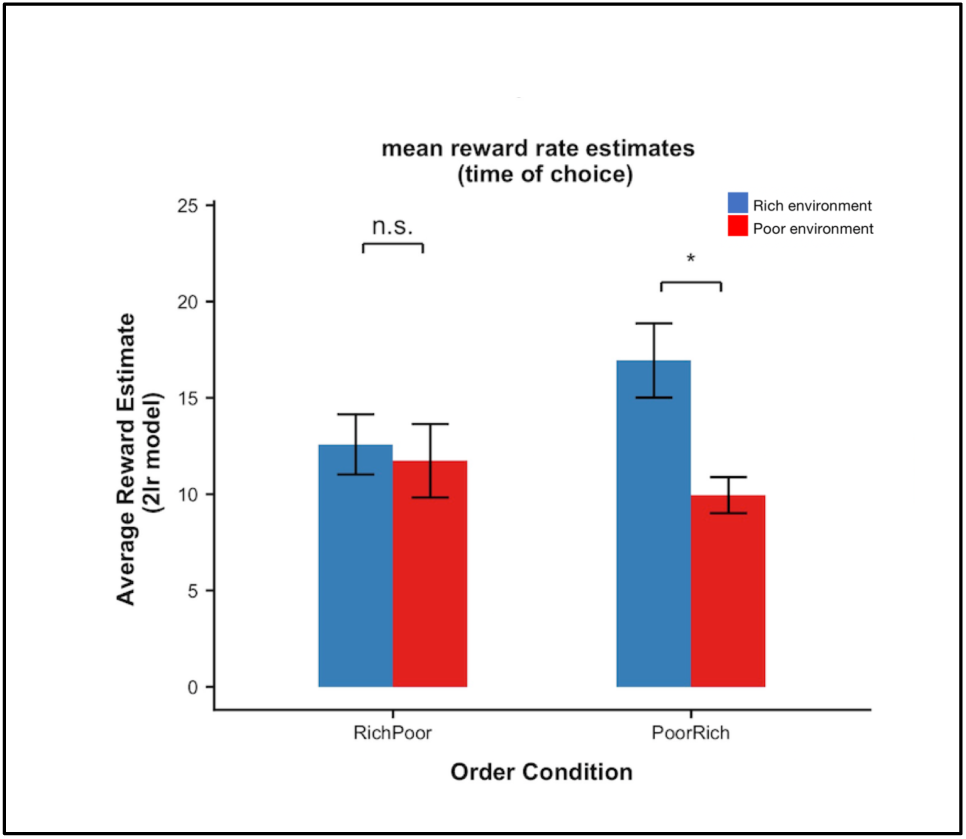
Consistent with Experiment 1 (main text and Figure 4a), extracting reward rate estimates (*ρ*) from the Asymmetry Model for each participant in each experimental block in Experiment 2 revealed a significant environment by condition interaction (F(1, 36) = 18.87, p<0.001). This arose out of a significant difference in *ρ* between environments for participants in the PoorRich condition (t(20) = 6.59, p<0.001, paired sample ttest) which was absent among participants assigned to the RichPoor condition (t(16) = 0.98, P=0.34). * p<0.05, paired sample ttest n.s. = non significant (p>0.05)

**Supplementary Figure 3.**
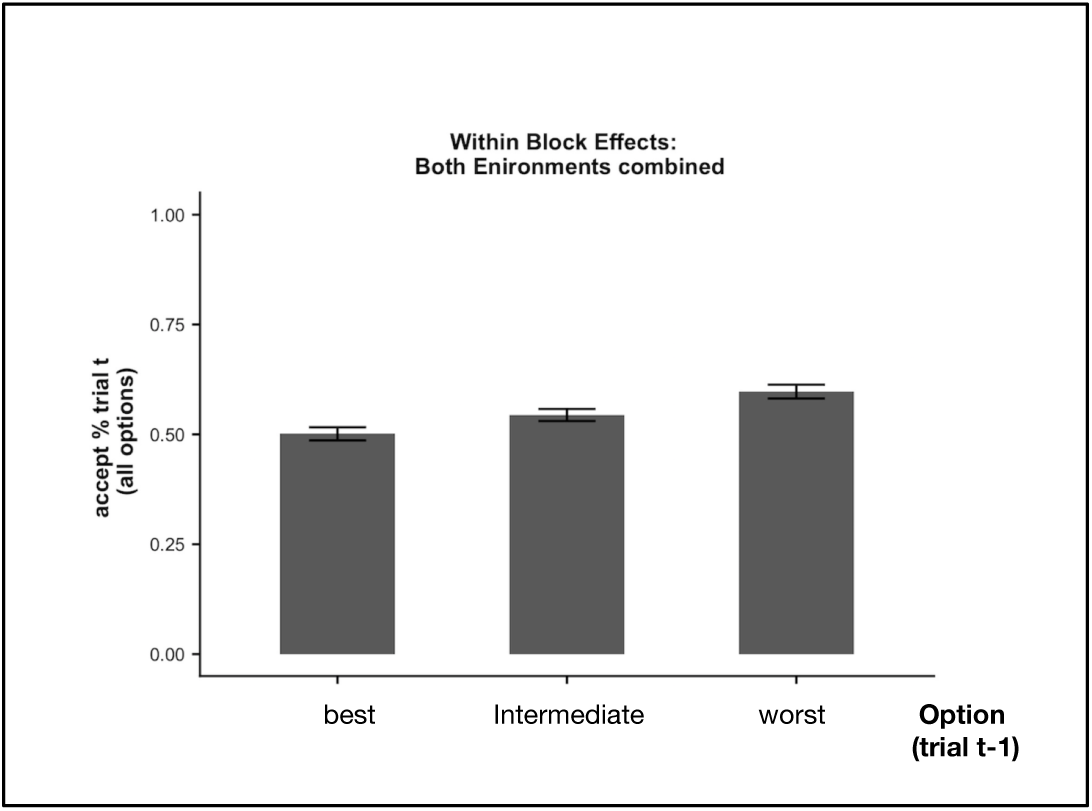
As observed in Experiments 1 and 2 (see main text and Figure 2), acceptance rates were modulated by trial to trial dynamics (main effect of previous option, controlling for environment: F(2, 74) = 21.02, p<0.001))

